# Targeting MOG to skin macrophages prevents EAE in macaques through TGFβ-induced peripheral tolerance

**DOI:** 10.1101/571828

**Authors:** Claire-Maëlle Fovet, Lev Stimmer, Vanessa Contreras, Philippe Horellou, Audrey Hubert, Nabila Sediki, Catherine Chapon, Sabine Tricot, Carole Leroy, Julien Flament, Julie Massonneau, Nicolas Tchitchek, Bert A. ’t Hart, Sandra Zurawski, Peter Klucar, Kumaran Deiva, Gerard Zurawski, SangKon Oh, Roger Le Grand, Ché Serguera

## Abstract

To study the effect of vaccination on tolerization to the myelin antigen MOG we used a macaque model of experimental autoimmune encephalitis (EAE) in which immunization with recombinant human myelin oligodendrocyte glycoprotein (rhMOG) elicits brain inflammation and demyelination mediated by MOG-specific autoreactive CD4+ T lymphocytes and anti-MOG IgG. For antigen targeting to tolerizing antigen presenting cells, we used a recombinant antibody directed to the Dendritic Cells (DC)-Asialoglycoprotein receptor (DC-ASGPR). The intradermal administration of an anti-DC-ASGPR-MOG fusion protein, but not a control anti-DC-ASGPR-PSA (prostate specific antigen) protein, protected monkeys committed to develop EAE. Although effective treatment did not modify anti-MOG IgG production, it prevented the CD4+ T lymphocyte activation and pro-inflammatory cytokine production. Moreover, animals treated with anti-DC-ASGPR-MOG experienced a rise of MOG-specific CD4+CD25+FOXP3+CD39+ regulatory T cells as well as a TGF*β*1, TGF*β*2 and IL-8 upsurge after rhMOG re-immunization. Our results indicate that the pathogenicity of autoantibodies directed to MOG is mitigated in the presence of MOG-specific regulatory lymphocytes. This vaccination scheme appears suitable to treat relapsing autoimmune diseases with identified autoantigens such as that harboring anti-MOG or anti-AQP4 autoantibodies.

## Introduction

Autoimmune demyelinating diseases (ADD) represent a major cause of non-traumatic neurological diseases in children and adults. The clinical spectrum is heterogeneous, evolving as a monophasic acute demyelinating encephalomyelitis, optic neuritis and transverse myelitis, or as a chronic disease such as Neuromyelitis Optica Spectrum Disorder (NMOSD) or multiple sclerosis (MS). Among these diseases NMOSD regroup rare subsets of autoimmune conditions for which self-targeted autoantigens have been identified as indicated by the high prevalence in patients of pathogenic IgG directed either to aquaporin 4 (AQP4) or to myelin oligodendrocyte glycoprotein (MOG) (Kitley *et al*, 2014). Due to the particular severity of these diseases an effective treatment is eagerly awaited and immune tolerization to AQP4 or to MOG could set a cure.

We have established a model of experimental autoimmune encephalitis (EAE) in cynomolgus macaques, which due to clinical-pathological and immunological similarities with ADD is exquisitely useful to investigate innovative therapeutic strategies for these diseases. The model is obtained through immunization with a recombinant protein representing the extracellular domain of human MOG, emulsified with the mineral oil IFA (Haanstra *et al*, 2013). MOG is a central nervous system (CNS) restricted myelin protein highly prevalent as a target of autoimmunity in ADD (Reindl *et al*, 2013); it is also a dominant antigen for chronic EAE induction in rodents and non-human primates (NHP) (t Hart *et al*, 2011). Cynomolgus macaques challenged with rhMOG/IFA develop an EAE characterized by neurological signs correlated with diagnostic brain lesions visible on MRI. Prominent immunological hallmarks are CD4+ T cells reactivity, and anti-MOG IgG. Histology shows that lesions detected with brain MRI correspond to demyelination with immunoglobulin deposits, complement activation and infiltration of neutrophils, macrophages as well as T and B-lymphocytes, testifying for a myelin-directed autoimmune aggression (Haanstra *et al*, 2013).

Pathogenesis of ADD, including EAE, presumes breakage of self-tolerance due to down-tuned regulatory T cells (Treg) and boosted activation of auto-reactive naïve or memory CD4+ T cells through MHC-II presentation of myelin or myelin-like antigen by specialized antigen presenting cells (APC) such as dendritic cells (DC). This leads to increased differentiation of CD4+ T lymphocytes into encephalitogenic Th1 and Th17 cells, either producing IFN-*γ* or IL-17, both favoring coupled infiltration of inflammatory macrophages and neutrophils into the brain white matter. In fact, studies in adults and children with MS or NMO have reported deficient Treg function (Brill *et al*, 2019) and abnormally increased Th1 and Th17 effector cells responses in the periphery (Kaneko *et al*, 2018; Sagan *et al*, 2016), which suggests that pathogenicity due to auto-reactive T cells could be counterbalanced by restoring proper Treg functions. This idea was corroborated in mouse models of EAE, either via adoptive transfer of Tregs *(Selvaraj & Geiger, 2008)* or through *in vivo* manipulation of DC for induction of MOG-specific Tregs (Idoyaga *et al*, 2013; Ring *et al*, 2013).

Dendritic cells (DCs) are the most potent antigen presenting cells (APCs) that can induce and direct adaptive response toward either immunity or tolerance *(Banchereau & Steinman, 1998)*. Hence, with the clinical purpose to control adaptive autoimmune response, DC-targeted vaccines are currently being developed (Palucka *et al*, 2010). Notably, subsets of immature migratory DCs from skin, gut or lungs have tolerogenic properties. In the absence of inflammation, they capture local antigens that they present to lymphocytes in draining lymph nodes, inducing their differentiation into antigen-specific Treg cells (Steinman *et al*, 2003). This is determined by particular co-stimulation of lymphocytes by DC secreting IL-10 and TGF-*β* (Selvaraj & Geiger, 2008; Ring *et al*, 2013; Li *et al*, 2012; Yamazaki *et al*, 2008).

In human skin, immature dermal DC, but not Langerhans cells express DC-asialoglycoprotein receptor (DC-ASGPR/CLEC10A), a C-type lectin scavenging receptor (CLR) that allows rapid endocytosis of ligands for antigen processing (Valladeau *et al*, 2001). We previously demonstrated that antigens (Ags) delivered to skin DCs via DC-ASGPR in macaques induce Ag-specific IL-10-producing CD4+ T cells testifying a regulatory function. In contrast, the targeting of the same Ag with anti-LOX-1 antibodies, induced IFN-γ producing T cell responses (Li *et al*, 2012).

Here, in a preclinical study in macaques immunized with rhMOG/IFA, we tested the clinical and biological effect of anti-DC-ASGPR-MOG immunotherapy on the induction and progression of EAE, a generic model of human autoimmune disease that is specifically projected on the autoimmune neuro-inflammatory disease MS. We report the effectiveness of anti-DC-ASGPR-MOG on the modulation of MOG-specific immune response.

## Results

### Clinical outcome of treatment with anti-DC-ASGPR-MOG

Anti-DC-ASGPR binds *in vitro* generated human monocyte-derived DC as well as CD11c+ and CD14+ cells in macaque PBMC (Valladeau *et al*, 2001). It also bind to CD14+CD1c+ dermal DCs in human skin (Li *et al*, 2012; Banchereau *et al*, 2012). To assess if anti-DC-ASGPR-MOG and rhMOG proteins target the same cells when injected in macaques’ dermis, we performed IHC on skin biopsies and observed that they respectively located either in CD163+ dermal macrophages or in CD1a+ cutaneous DC (**figure 1A and 1B**) (Zaba *et al*, 2007; Adam *et al*, 2015). Moreover, rhMOG, but not the anti-DC-ASGPR-MOG, was associated with the expression of CD40 (**fig. S1**). This indicates that the anti-DC-ASGPR-MOG can be captured by CD163+CD40-resident macrophages in an anti-DC-ASGPR mAb-specific manner, while rhMOG is phagocytosed by CD1a+ CD40+ Langerhans cells or dermal DCs.

**Figure 1:**
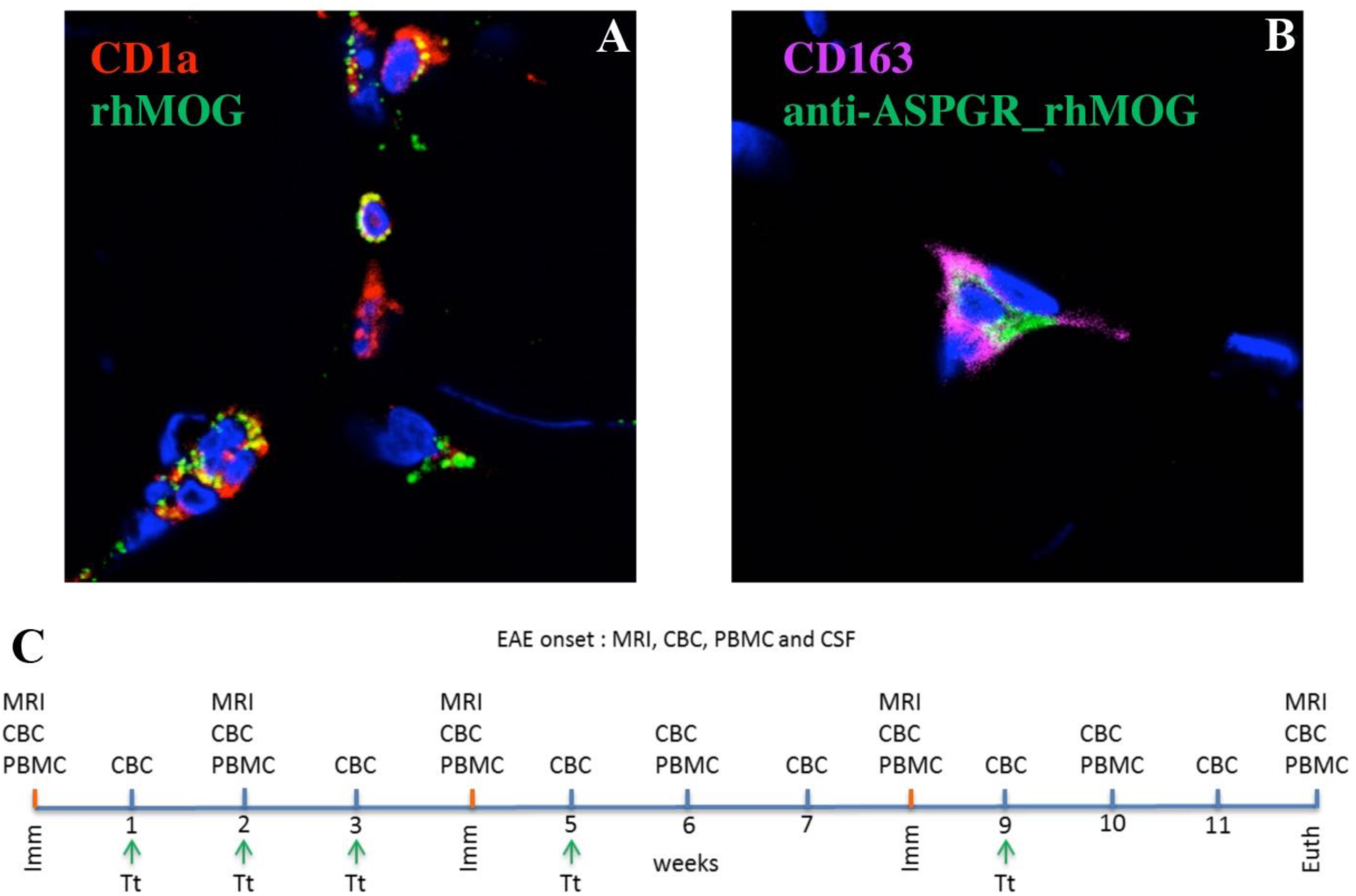
*Localization of injected proteins to macaques in vivo and experimental design.* rhMOG or anti-DC-ASGPR-MOG conjugated to AF488 were injected intradermally and a skin biopsy of the injection site was analyzed 4hr later with IHC. **A**) rhMOG was detected in CD1a+ cells close to the epidermis. **B**) anti-DC-ASGPR-MOG colocalized with CD163+ dermal cells. **C**) Preclinical experiments were carried out over 12 weeks. Cynomolgus macaques were immunized with rhMOG/IFA every 4 weeks (red ticks), until onset of EAE. Treatment with anti-DC-ASGPR-MOG or anti-DC-ASGPR-PSA (green arrows) was administered every week for 3 weeks after initial immunization and then every week after the boost with rhMOG/IFA. MRI and PBMC collection were undertaken at day 0 and then every 2 weeks. CBC was performed every week. CSF was collected at EAE onset. Abbreviations: Imm, immunization; MRI, magnetic resonance imaging; CBC, complete blood count; CSF, cerebrospinal fluid; Euth, euthanasia. Tt, treatment; IHC immunohistochemistry. Nuclei were stained with 4,6 diamidino-2, phenylindole (DAPI).

In cynomolgus macaques rhMOG/IFA immunization leads to clinically evident EAE in about 35 days (Haanstra *et al*, 2013). We treated rhMOG-sensitized monkeys (n=6) with anti-DC-ASGPR-MOG (n=3) or an anti-DC-ASGPR-PSA (n=3). Animals were followed as detailed in (**figure 1C**). The 3 animals receiving the control anti-DC-ASGPR-PSA fusion protein developed clinical signs of EAE. Two of them (C1 and C2), developed a classical EAE with onset at 22 and 32 dpi, evolving towards a severe disease in 14 and 3 days (**figure 2A**). The third control animal faced several episodes of paresthesia and tremor with lower clinical score between 24 and 30 dpi (animal C3, **figure 2A**). On the contrary, none of the 3 macaques treated with anti-DC-ASGPR-MOG developed clinical signs of EAE within 90 dpi (**figure 2B**). This divergent response to rhMOG immunization points to a therapeutic effect of the treatment with the anti-DC-ASGPR-MOG.

**Figure 2:**
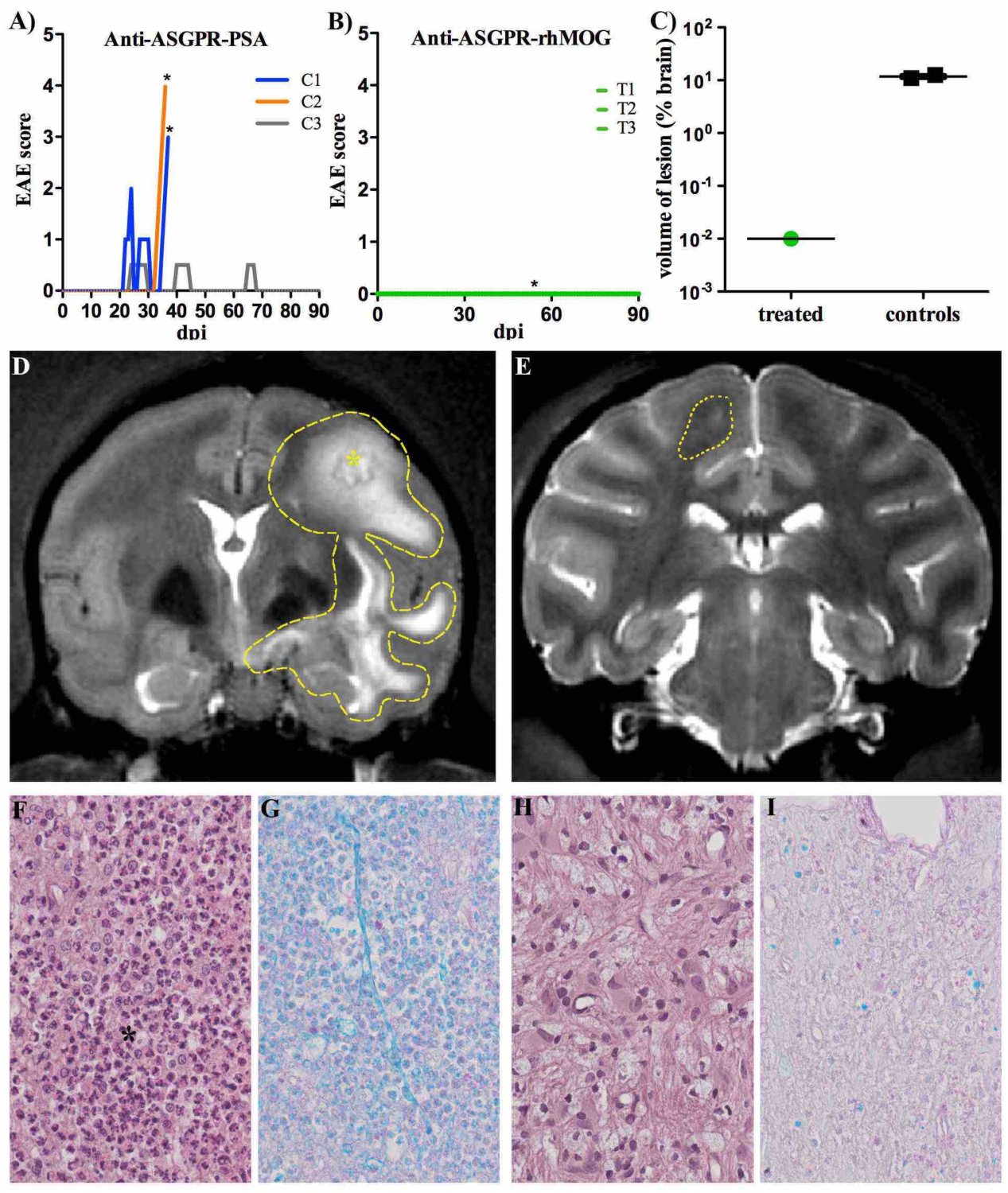
*Clinical features, imaging and histology of EAE in treated and control animals.* Onset of EAE was detected through clinical observation of animals. Behavioral and neurological deficits were rated according to a grid allowing scoring the severity of disease (**supplementary table 2**). **A)** Onset, severity and progression of EAE in control animals (C1, C2 and C3) from day 1 to 90 post-immunization (dpi); higher scores reflecting deeper illness. Stars (*) represent brain lesions detected with MRI. **B)** onset and progression of disease in treated animals. Green lines represent basal scoring in treated animals (T1, T2 and T3). **C)** Volume of brain lesions in control and treated animals measured on MRI. Note a 10^3^x higher volume of lesions in controls as compared to the treated animal. Two pictures of MRI with, **D)** a characteristic EAE lesion imaged in animal C2 at 35 dpi circled with a dot line, and **E)** a small lesion detected at 54 dpi in animal T2 circled with dote line. **F-I)**: Histopathology of WM lesions in control (**F and G**) and treated (**H and I**) animals, magnification 400x; (**F and H**) Hematoxylin Eosin (HE) staining; (**G and I**) Luxol-Fast-Blue-PAS (LFB) stain for myelin fibers; Control animals (**F**) show severe myelin degeneration with an infiltration of a high number of neutrophils (black star) and low number of activated microglia/macrophages (black arrow), whereas the lesion found in one treated animal (**H**) displays low cellularity infiltrated mainly by few activated microglia/macrophages (black arrows). Severe myelin phagocytosis is present in lesions of control animals (**G**) shown by numerous LFB positive granules in inflammatory cells, whereas only a low amount of myelin is present in macrophages of the lesion of the treated animal (**I**) indicating a chronic character of this lesion.

In line with clinical observations, the 2 control animals that developed severe EAE, extended hyperintense signals that were detected with MRI, indicating brain inflammation (**figure 2D**). In one animal treated with anti-DC-ASGPR-MOG, a lesion was also detected with MRI at 54 dpi (**figure 2E)**, but it was about 1000 times smaller than those observed in control animals **(figure 2C**) and was never associated to any clinical sign of EAE (**figure 2B**). This lesion was transient as it had disappeared at a later MRI (82 dpi).

For histology, brain lesions detected with MRI, were dissected; these lesions presented different histopathology in 1 treated and 2 control animals (**fig. S2**). Brains of control animals developing a severe EAE, each presented 2 and 3 confluent large lesions of destroyed white matter and some adjacent gray matter. These were round, confluent and often centered on vascular structures surrounded by necrotic WM with hemorrhages and infiltrated by high numbers of degenerated neutrophils and vacuolated macrophages (**figure 2F**). Luxol fast blue (LFB) special stain for myelin showed demyelination involving virtually the entire lesion observed in HE (**figure 2G**). Furthermore, vacuolated macrophages contained a high amount of phagocytosed LFB-positive myelin debris, confirming an active demyelination. As observed with MRI, treated animal T2 presented a small subcortical lesion. It contained only few vacuolated macrophages and no neutrophils (**figure 2H**). LFB special stain showed moderate demyelination. However, most macrophages did not contain LFB positive myelin debris, reflecting a chronic inactive lesion (**figure 2I**). Analysis of the brain and spine of the two other treated animals revealed no lesion. Control animal C3 developed a mild EAE with signs evoking a spinal-cord lesion. However, our MRI setting does not allow surveying the spinal cord. As we missed precise coordinates, we were not able to detect the lesion.

### MOG-specific antibody response

Anti-MOG immunoglobulins are a pathologically relevant feature in our EAE model (Haanstra *et al*, 2013). ELISA revealed that all animals experienced a rapid surge of anti-MOG IgG following rhMOG/IFA immunization, with no differences between treated and control animals (**figure 3A**). As only antibodies binding conformationally intact epitopes of native MOG can be pathogenic, we titrated anti-MOG-IgG1 in a cell-based assay (CBA). Binding to cellular MOG by plasmatic IgG1 displayed individual trends but was not consistently different between groups. Animal C1 developing a progressive form of EAE had intermediate levels of anti-MOG IgG1. Animal C2 presenting an abrupt EAE had elevated anti-MOG IgG1 and animal C3 with mild EAE had the lowest titer of anti-MOG IgG1. These observations are consistent with the severity of disease being correlated to the level of anti-MOG IgG1. However, treated animals T1 and T2 also had high levels of anti-MOG IgG1 at 35 dpi and thereafter, but resolutely remained asymptomatic (**figure 3B**). This indicates that the production of anti-MOG immunoglobulin was not modulated by anti-DC-ASGPR-MOG and that high levels of circulating anti-MOG IgG1 is not sufficient to trigger clinical EAE. This points out that the pathogenic factor suppressed by the treatment is not related to anti-MOG IgG1.

**Figure 3:**
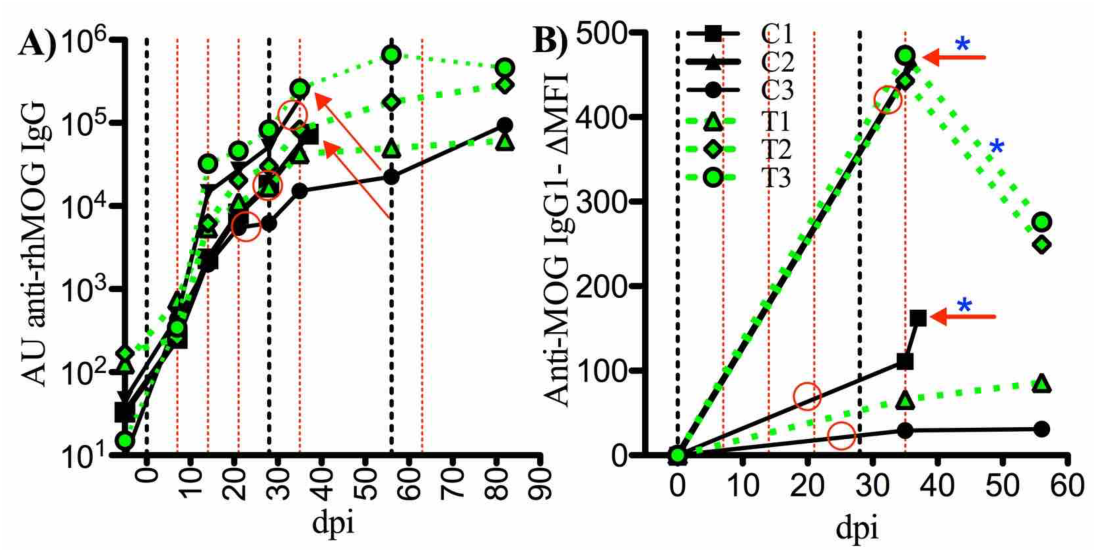
*Plasma levels of anti-rhMOG IgG measured with ELISA and CBA.* Levels of IgG binding to plate-bound rhMOG in plasma of animals were measured every week with ELISA. **A)** Levels of arbitrary units (AU) of anti rhMOG IgG increase for several weeks. **B)** anti-MOG IgG1 measured with cell-based assay (CBA) at baseline and at 35 dpi for all animals as well as at 36 dpi for animal C1 and 37 dpi for animal C2; levels of IgG1 are expressed in *Δ*MFI (see methods). Control animals: black lines; treated animals: green lines. Vertical dotted black lines: rhMOG/IFA immunizations, vertical dotted red lines: anti-DC-ASGPR-MOG or -PSA injection; red circles: EAE onset, red arrowheads: euthanasia and blue stars: a brain lesion detected with MRI.

### Increased neutrophils and drops of lymphocytes at disease onset

We had previously noticed that EAE onset and progression in cynomolgus macaques are correlated with a dramatic surge of circulating neutrophils and a concomitant drop of lymphocytes resulting in a rising neutrophil to lymphocytes ratio (NLR) (our unpublished results). Here, we also observed that severe EAE in control animals was associated with an abrupt increase of circulating neutrophils and a concomitant drop of circulating lymphocytes, starting at onset of disease leading to a steep NLR increase (C1 and C2, **fig. S2**). In the control animal C3 with milder EAE the transient clinical signs detected at 24-29 dpi was also associated to weak peaks of NLR. In the 3 animals treated with the anti-DC-ASGPR-MOG, although 1 of them had a small cortical lesion at MRI, blood leukocytes remained at base level suggesting that the treatment prevented neutrophils outburst and lymphocytes decline in the blood (**fig. S2**). These findings point to a pathogenic role of neutrophils, as also noticed in rhesus macaque EAE (Dunham *et al*, 2017).

### Phenotype of Circulating Lymphocytes

Characterization of circulating lymphocytes revealed that the percentage of CD4+ T cells tended to increase in treated animals after immunization with rhMOG (**figure 4A**), but that activated CD4+ T cells (CD69+) were significantly increased only in controls at 30 dpi as compared to earlier time points (p<0.05) (**figure 4B**). Moreover, activated central memory (CM) CD4+CD95+CD28+CD69+ T lymphocytes were also increased at 30 dpi in controls as compared to treated animals (p<0.05) (**figure 4D**).

**Figure 4:**
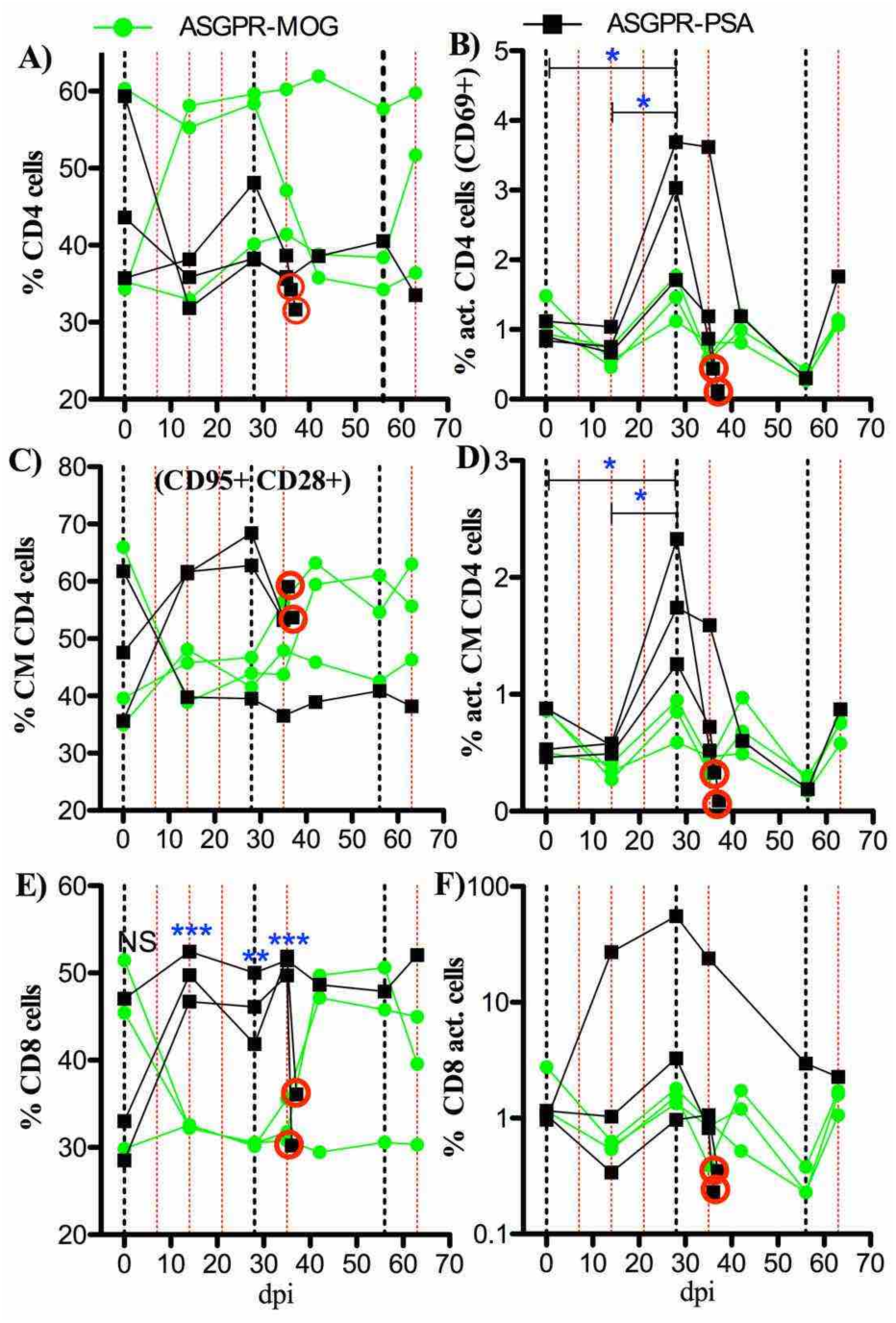
*T cell phenotypes.* Phenotype of circulating CD4+ and CD8+ T lymphocytes assessed with flow cytometry at several time points after immunization. A) treated animals tend to have more circulating CD4+ T cells than controls. B) activated CD4+CD69+ cells are increased in controls at 35 dpi as compared to earlier time points. C) Central memory CD4+CD95+CD28+ cells tended to increase in control animals after immunization with rhMOG/IFA. D) activated central memory CD4+ CD95+CD28+CD69+ cells were significantly increased in control animals at 28 dpi. E) control animals have increased levels of circulating CD8+ T cells compared to those observed in treated animals. F) activated CD8+CD69+ cells tended to increase in the control animal developing a mild EAE. Curves represents follow-up of each animal, in green for those treated with anti-DC-ASGPR-MOG and in black for control animals treated with anti-DC-ASGPR-PSA. Red circles: euthanasia at pick of EAE severity. Statistics, unpaired t-test, two-tailed; (*) p< 0.05; (**) p< 0.01; (***) p < 0.001.

CD8+ T cells increased significantly in control animals after immunization with rhMOG from 14 to 35 dpi, but not in treated animals (p<0.001) (**figure 4E**). This was associated with an increase of activated CD8+ only for control animal (C3) developing a mild EAE (**figures 4F**) suggesting a particular pathological progression.

No difference appeared in circulating NK nor B cells naïve or memory subtypes whether activated or not (not shown).

Thus, it appears that the treatment with anti-DC-ASGPR-MOG prevented activation of CD4+ lymphocytes as well as proliferation of circulating CD8+ lymphocytes following rhMOG immunization.

### Cytokine levels in plasma

We measured plasma levels of 15 cytokines at different time points in treated and control monkeys and in 4 naïve animals. A heatmap revealed 4 groups, which from right to left are ordered from lower to higher content of cytokines (**figure 5A and fig. S4**); Group I encloses only treated animals at 21 and 28 dpi depicting a “disease solving” profile with particularly low levels of TGF*β*2 and higher levels of IL-10. Group II, is the most heterogeneous and includes all treated and control animals at 7 dpi, as well as some animals of either group at 21 or 28 dpi but no naïve animals, pointing at a “disease incubation” pattern with a significant decrease of TGF*β*1 (p=0.0002), TGF*β*2 (p=0.00002) and IL8 (p=0.01) as compared to naïve animals (**Figure 5B**). Group III aggregates the naïve macaques, the 3 treated animals remaining healthy at 35 dpi (1 week after 1^st^ boost with rhMOG/IFA) and control C3 at 35 dpi, in remission from a mild EAE. This group then relates to a “healthy pattern” of cytokines with “restored” levels of IL8, TGF*β*1 and TGF*β*2, and treated macaques displaying even higher levels of TGF*β*1 (p=0.002), TGF*β*2 (p=0.006) and IL8 (p=0.03) than naïve animals (**Figure 5C**). Group IV gathers only animals treated with anti-DC-ASGPR-PSA at onset or peak of EAE with decreased TGF*β*1 (p=0.008), TGF*β*2 (p=0.015) but not IL8 (p=0.05) as compared to naïve animals and with concomitant increased levels of pro-inflammatory cytokines (see below). These comparisons reveal that EAE incubation and onset are associated to a drop of systemic levels of TGF*β*1, TGF*β*2 and IL8 as well as to an increase in pro-inflammatory cytokines, and that the treatment restores TGF*β*1, TGF*β*2 and IL8 expression to physiological levels or above and prevents burst of pro-inflammatory cytokines.

**Figure 5:**
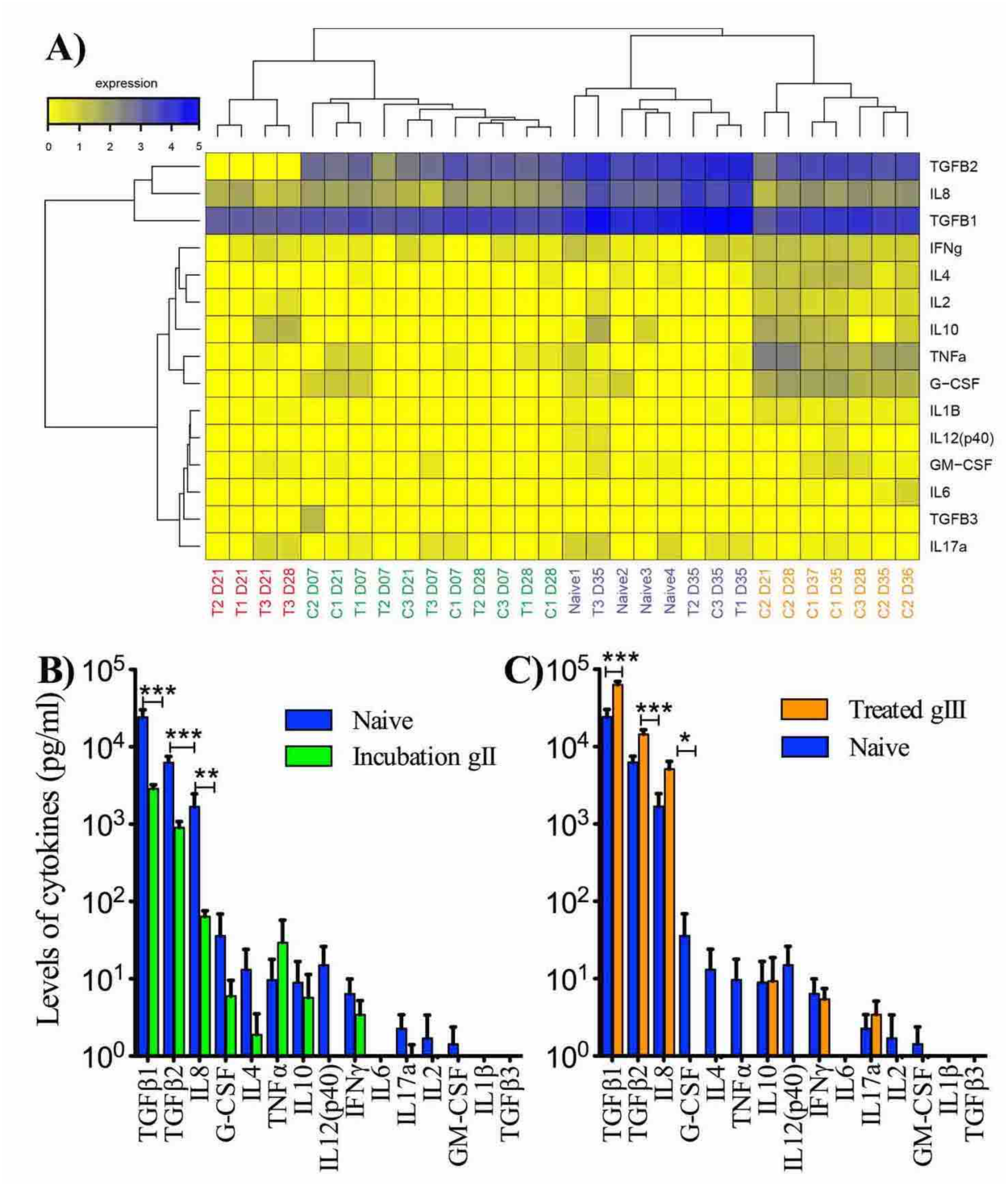
*Cytokine dosage in plasma of treated and control animals*. **A)** Heatmap showing cytokine levels in plasma for all animals at several time points after immunization with rhMOG/IFA. Expression levels of cytokines are represented with a color gradient ranging from yellow (no expression) to deep blue (highest concentration). Concentrations are expressed in log10 of pg/ml. Hierarchical clustering, represented by dendrograms were performed at individual and cytokines levels. Four groups of animals were identified, having low to high amounts of cytokines and numbered from 1 to 4 in the text; animals treated with anti-DC-ASGPR-MOG are named (T1, T2, T3), controls treated with anti-DC-ASGPR-PSA are named (C1, C2, C3), and untreated non-immunized animals are named (naïve 1, 2, 3, 4); time points in days post-immunization are numbered from D7 to D37; group 1: red (EAE solving), group 2: green (EAE incubation), group 3: purple (EAE resolution), group 4: yellow (EAE onset and progression). See also **table S4**. **B)** Cytokine levels in naïve animals as compared to that measured in group 2 referred as EAE incubation group where a significant decrease in TGF*β*1, TGF*β*2 and IL8 is observed. **C)** Cytokine levels in naïve animals as compared to that measured in treated animals of group 3 where a significant increase in TGF*β*1, TGF*β*2 and IL8 is observed. Statistics, unpaired t-test, two-tailed; (ns) p > 0.05; (*) p< 0.05; (**) p< 0.01; (***) p < 0.001.

Statistical comparison of cytokine levels between individuals at different times indicates increasing amounts of the pro-inflammatory cytokines IFN*γ*, IL1*β*, G-CSF, GM-CSF or TNF*α* in controls, while treated animals rather experienced a significant increase in IL8, TGF*β*1 and TGF*β*2 after reimmunization at 35 dpi (**fig. S3**). In fact, comparing cytokine levels between the two groups shows that the 3 controls, but not treated animals, had increased levels of the pro-inflammatory cytokines IL-1*β* (p=0.01), IFN*γ* (p=6×10^−4^) and TNF*α* (p=0.01) at disease onset (35 dpi) as compared to earlier time points. Instead, TGF*β*1 (p=3×10^−7^) TGF*β*2 (p=3×10^−5^) and IL-8 (p=9×10^−5^), were notably increased in treated animals at 35 dpi as compared to earlier time points, which was not the case in controls (**figure 6**).

**Figure 6:**
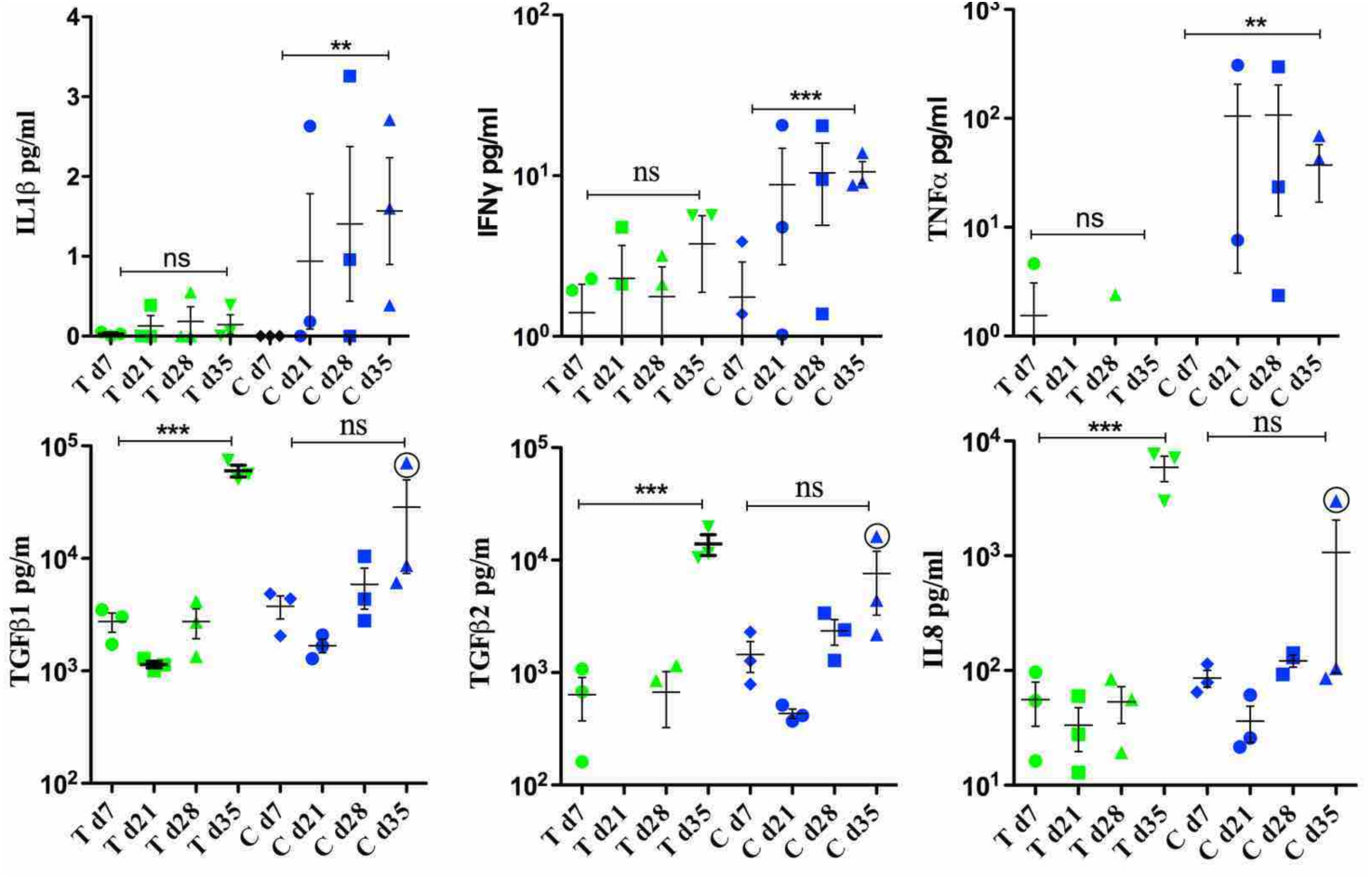
*Levels of several cytokines at different time points after immunization with rhMOG/IFA.* In green, animals treated with anti-DC-ASGPR-MOG (T); in black, control animals treated with anti-DC-ASGPR-PSA (C). Pro-inflammatory cytokines IL1*β*, IFN*γ*, or TNF*α* are elevated in controls but not in treated animals at latest time point of 35 dpi. Levels of IL8, TGF*β*1 and TGF*β*2 are elevated in treated animals at latest time point of 35 dpi, but not in controls. In one animal treated with anti-DC-ASGPR-PSA (C3) developing a mild EAE between 24 and 29 dpi (circled), we also observe higher levels of TGF*β*1, TGF*β*2 and IL8 at 35 dpi that are significantly higher than that measured in C1 and C2 developing a severe EAE at this precise time point (see supplementary figure 3). Statistics, unpaired t-test, two-tailed; (ns) p > 0.05; (*) p< 0.05; (**) p< 0.01; (***) p < 0.001.

Interestingly, at 28 and 35 dpi, the resilient animal C3, which had earlier developed a mild form of EAE, displayed a peculiar intermediary pattern of cytokines to that measured in animals with full-blown EAE and in treated animals. It combined both patterns of pro-and anti-inflammatory cytokines, suggesting that at some point the latter were sufficient to control the disease progression (**fig. S3**).

### MOG-specific regulatory cells

TGF-*β*1 prevents spontaneous activation of naïve CD4+ T cells and the induction of widespread autoimmunity (Gorelik & Flavell, 2000; Tu *et al*, 2018) in part through induction of peripheral FOXP3+ Treg lymphocytes (Chen *et al*, 2003). Thus, taking advantage of a sensitive test measuring lymphocyte activation through CD25 and CD134 (OX40) expression shortly after specific-antigen stimulation (Zaunders *et al*, 2009), we assessed whether the increase in blood TGF*β* in treated animals was associated with an increase in MOG-specific CD4+CD25+OX40+FOXP3+ T cells. Moreover, by measuring the surface expression of the ectoenzyme CD39, we asked if these cells originated from resting memory CD39+ Tregs, a subset of cells with suppressor properties and stable expression of FOXP3 (Seddiki *et al*, 2014). In PBMC stimulated with rhMOG, MOG-specific CD4+CD25+OX40+ remained steady in both groups (**Figure 7A**). Among these, rates of FOXP3+CD39-cells also remained stable in treated and controls (**Figure 7B**), while MOG-specific FOXP3+CD39+ lymphocytes were increased at 35 dpi in treated animals but not in controls (**Figure 7C**).

**Figure 7:**
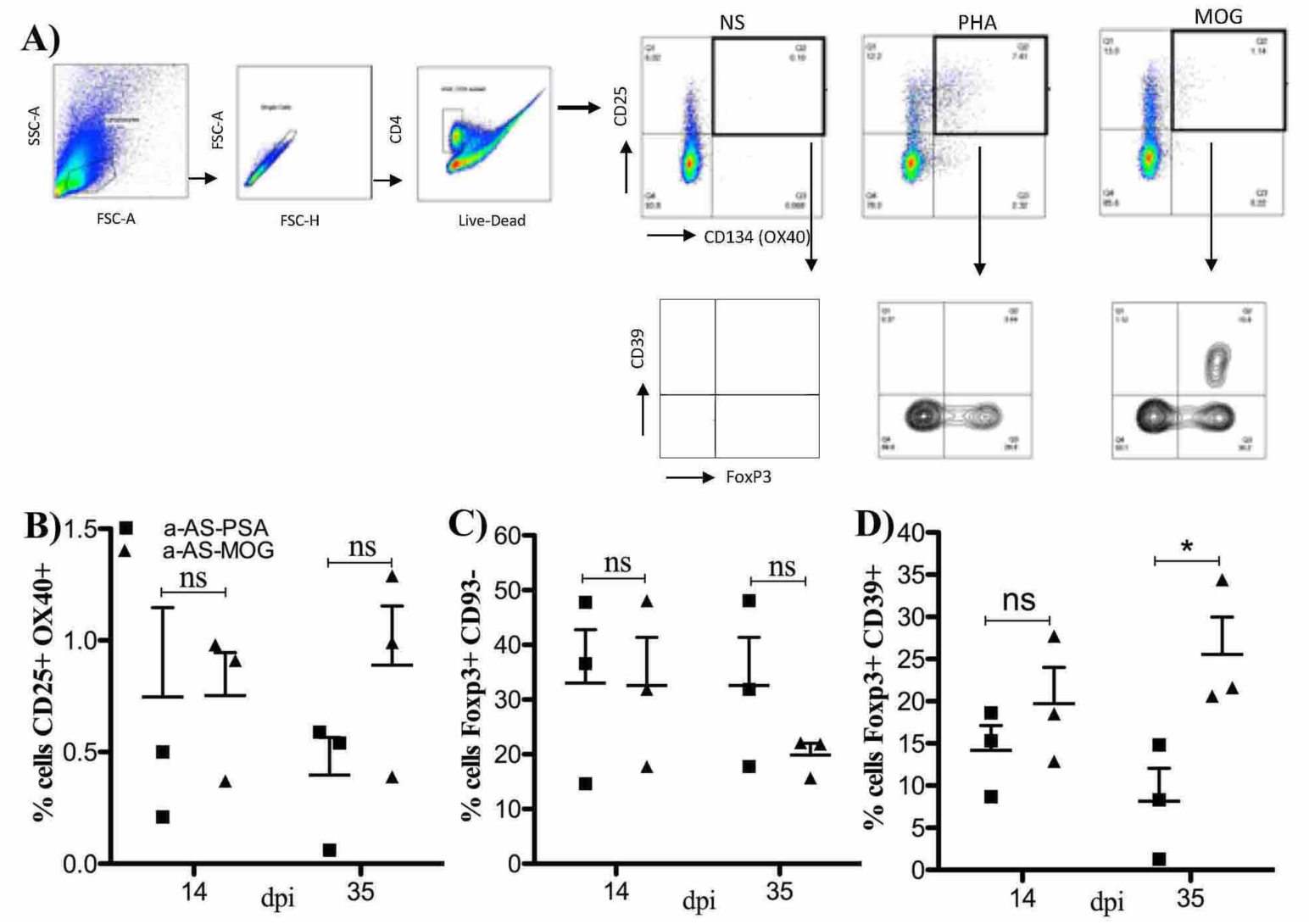
*Treg lymphocytes measurments.* We determined FOXP3 and CD39 expression among CD4+CD25+CD134+ T cells Ag recall response in macaques immunized with rhMOG/IFA and receiving anti-DC-ASGPR-MOG (a-AS-MOG) or anti-DC-ASGPR-PSA (a-AS-PSA). **A)** PBMC from macaques at 14 or 35 dpi, were either left unstimulated (NS) or stimulated with PHA or rhMOG (MOG) for 44 hr, and MOG-specific responses analyzed by flow cytometry. The upper right quadrant shows CD4+ T cells responding to each stimulus identified as CD25+CD134+ (top plots). In lower plots, we observed the percent of each specific CD4+CD25+CD134+ T cell response that were also CD39+ (right upper quadrant) or FOXP3+ (right lower quadrant). **B)** Percent of CD4+CD25+CD134+ T-cells in PBMC of animals treated (a-AS-MOG) or controls (a-AS-PSA) at 14 or 35 dpi. **C)** Percent of FOXP3+CD39-among CD4+CD25+CD134+ T-cells in treated and control animals. **D)** Percent of FOXP3+CD39+ among CD4+CD25+CD134+ T-cells in in treated and control animals. Statistics, unpaired t-test, two-tailed; (ns) p > 0.05; (*) p< 0.05.

### Preventive vaccination with anti-DC-ASGPR-MOG

To assess if anti-DC-ASGPR-MOG alone induces anti-MOG IgG, 2 macaques were treated with this antibody-antigen chimeric protein and then challenged with rhMOG/IFA. Anti-MOG IgG remained at basal levels during treatment with anti-DC-ASGPR-MOG and increased only after rhMOG/IFA immunization, indicating that although fused to the exact same protein sequence as rhMOG, the anti-DC-ASGPR-MOG does not stimulate anti-MOG IgG production. Instead, the treatment with anti-DC-ASGPR-MOG induced progressive increase of MOG-specific CD4+CD25+FOXP3+CD39+ cells, which were possibly recruited from a pool of memory cells or alternatively converted from more frequent circulating MOG-specific CD4+FOXP3+CD39-cells (**Figure 8B**). Finally, in spite of 3 successive immunization with rhMOG/IFA, these 2 animals receiving a preventive treatment remained healthy and expressed no sign of brain inflammation at MRI during at least 90 days following a last injection of anti-DC-ASGPR-MOG.

**Figure 8:**
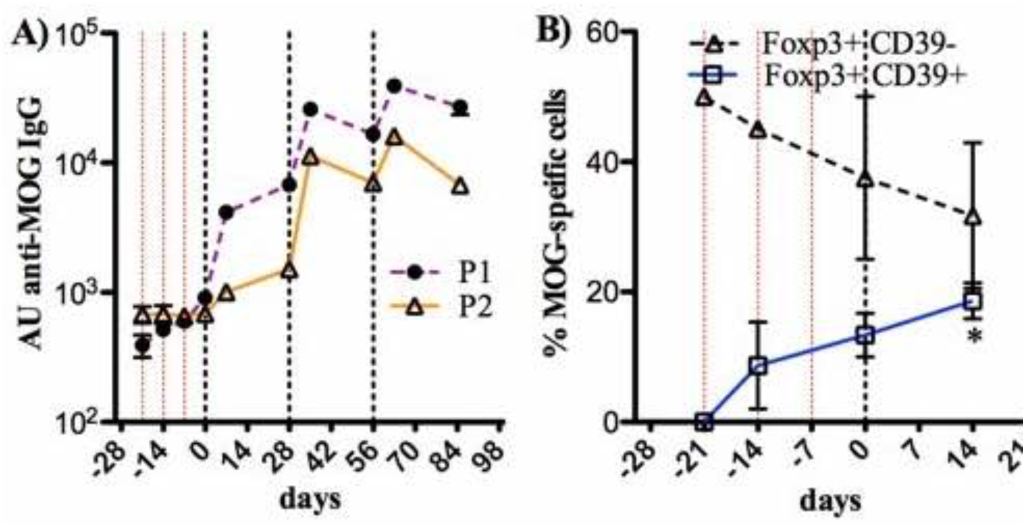
*Preventive vaccination with anti-DC-ASGPR-MOG.* Presence of anti-MOG IgG and CD4+CD25+CD134+ FOXP3+ CD39+ T cells in 2 macaques preventively treated with anti-DC-ASGPR-MOG. **A)** Weekly ELISA dosage of arbitrary units (AU) of anti-rhMOG IgG in plasma of animals. Anti-MOG IgG remained at basal levels during treatment with anti-DC-ASGPR-MOG but increase upon immunization with rhMOG/IFA. **B)** Percentage of CD4+CD25+CD134+ T cells that express FOXP3+CD39-(doted black line) or FOXP3+CD39+ T-cells (plain blue line) among PBMC of the 2 macaques at successive time points. In both graphs, thin dotted red lines: anti-DC-ASGPR-MOG injection; dotted black lines: rhMOG/IFA immunizations. Statistics, unpaired t-test, two-tailed; (*) p< 0.05.

## Discussion

Autoimmune conditions are commonly treated with immunosuppression while awaiting adapted therapies able to restore immune homeostasis and health. Here, we report an effective immune tolerization scheme blocking brain anti-MOG autoimmune burst in macaques. The cure based on dermal shipping of the myelin Ag MOG_1-125_ into resident skin macrophages favors the appearance of MOG-specific Tregs and a dramatic systemic surge of TGF*β* and IL-8 upon rhMOG/IFA re-stimulation.

The present results establish that a tolerogenic vaccine able to mitigate EAE in inbred SPF mice through MOG-targeting to peripheral DC and induction of FOXP3+ suppressor Tregs (Idoyaga *et al*, 2013; Ring *et al*, 2013) also applies to outbred adult primates with a trained immune system. In mice, tolerogenic priming relied on the ability of particular DC subsets to produce TGF*β* (Idoyaga *et al*, 2013; Ring *et al*, 2013; Yamazaki *et al*, 2008), a cytokine also produced by CD163+ M2 macrophages (Zaba *et al*, 2007; Alvarado-Vazquez *et al*, 2017). These CD163+ macaque’s skin macrophage have migratory properties as they are also retrieved in lymph nodes (Adam *et al*, 2015). In cancer and persistent mycobacteria infection, CD163+ M2 macrophages produce TGF*β* establishing an immunosuppressive environment inducing regulatory lymphocytes and subverting adaptive immune response (Fujimura *et al*, 2018; Yang *et al*, 2016). Thus, our results align with the idea that in physiological conditions these macaque CD163+ dermal resident macrophages are committed to the maintenance of tolerance to phagocytosed self-antigens.

Different from anti-DC-ASGPR-MOG, the intradermal injection of rhMOG was captured by skin CD1a+CD40+ DC, with pro-inflammatory properties (Salabert *et al*, 2016). Thus, animals receiving injections of both rhMOG and anti-DC-ASGPR-MOG, likely experienced concomitant priming of MOG-specific effector and regulatory lymphocytes with notorious restraint of otherwise strongly pathogenic autoreactive T cells. This was evidenced by an absence of disease and the lack of T cell activation and pro-inflammatory cytokines in the blood of treated animals contrary to what observed in controls.

Remarkably, all animals immunized with rhMOG/IFA went through a disease incubation pattern characterized by decreased plasma levels of TGF*β*1 and TGF*β*2. Current views associate EAE development to rising production of Th1 and Th17 cytokines (t Hart *et al*, 2011) while, to our knowledge, a systemic drop of anti-inflammatory cytokines in this context has not been described. Nonetheless, this point to a general homeostatic mechanism allowing naïve T cells priming prior to effector response. In fact, sustained TGF*β*1 signaling maintains naïve T cells quiescent and lack of this cytokine signaling in T cells causes widespread autoimmune Th1 response (Gorelik & Flavell, 2000; Tu *et al*, 2018). Moreover, a fall of plasma TGF*β*1 was observed at the start of Guillain-Barré, during MS relapse, in systemic lupus erythematosus or associated to effective immunization routes, suggesting that adaptive (auto)immune response is enabled by antigen presentation and co-stimulation but also by an environment lowering threshold for T cell activation (Bhowmick *et al*, 2009; Fletcher *et al*, 2008; Manolova *et al*, 2013; Rieckmann *et al*, 1994).

Interestingly, plasma levels of IL-8 mirrored that of TGF*β*1 and TGF*β*2 as it dropped during EAE incubation in all animals but increased dramatically after boost with rhMOG in all animals treated with anti-DC-ASGPR-MOG. This suggests that IL8 and TGF*β* may act in concert to maintain immune homeostasis. Numerous experiments establish the central role of TGF*β* in preserving immune homeostasis keeping naïve T cells quiescent and inducing their differentiation into regulatory or suppressor lymphocytes or restraining effectors (Selvaraj & Geiger, 2008; Ring *et al*, 2013; Yamazaki *et al*, 2008; Tu *et al*, 2018; Chen *et al*, 2003). On the contrary IL8 is commonly described as a major pro-inflammatory chemokine *(Paul, 2013)*, while it has also been implicated in homeostasis of subsets of naïve and regulatory CD4+ T lymphocytes (Crespo *et al*, 2018; Gibbons *et al*, 2014; Himmel *et al*, 2011). The reason why some regulatory T cells express IL8 is not understood, but its release by tumors, which can be induced by TGF*β* (Langhans *et al*, 2013; Bhola *et al*, 2013), promotes recruitment of Tregs as a mechanism of tumor escape from immune rejection (Eikawa *et al*, 2010). Another interesting phenomenon concerns a viral orthologue of IL8 expressed by Marek’s Disease Virus that favors virulence by attracting CD4+ CD25+ Tregs where the virus establishes latency and promotes lymphomagenesis (Engel *et al*, 2012). Thus IL8, a major chemokine in neutrophil activation and recruitment, seems to also attract regulatory CD4+ T cells to sites of inflammation to prevent indiscriminate activation of autoreactive lymphocytes. In our experiments MOG-specific regulatory CD4+ T lymphocytes likely responded to the reinjected rhMOG by producing IL8 and TGF*β* in proportion, then attracting neutrophils for antigen clearance but simultaneously preventing adapted response to an identified self-antigen (MOG). A similar sequence of events may take place at each encounter between MOG-specific Tregs and macaque’s own MOG epitopes presented by APC at the blood-brain-barrier, in brain draining lymph nodes or in contact to myelin, preventing effector T cells EAE induction (Spadaro *et al*, 2018).

Most interestingly a control animal having developed an early transient and low score EAE during the disease incubation phase, also produced high blood levels of TGF*β* and IL-8 after re-immunization with rhMOG, proving resilience to further progress of disease. Thus, the observed TGF*β*/IL-8 surge in response to an antigen likely corresponds to a physiological pattern of peripheral tolerance that can be reliably recapitulated and amplified by the treatment with anti-DC-ASGPR-Ag.

We had previously shown that the fusion of an antigen to anti-DC-ASGPR induced IL-10 secretion by human monocyte-derived DC and the priming of naïve CD4+ T cells into Ag-specific IL-10+ FOXP3-suppressors. When injected into macaque skin, this same fusion protein also favored the appearance of Ag-specific T cells secreting IL10, but their precise phenotype had not been assessed (Li *et al*, 2012). Here we show that skin injection of the anti-DC-ASGPR antibody induces MOG-specific CD4+ FOXP3+CD39+ Tregs. It is possible that MOG-specific IL10+FOXP3-suppressors had also been induced at particular time points of the protocol but due to difficulties in labeling IL10 in macaques’ PBMC we were not able to detect them. In addition, whether DC-ASGPR targeting into cultured IFNDC or in CD163+ dermal macrophages take alternative paths, needs to be addressed in future studies.

In conclusion, we report a preclinical protocol based on the vaccination with anti-DC-ASGPR-MOG, which induces robust protection of NHP against a grave tissue-specific autoimmune disease. The same approach might be applied to treat autoimmune diseases with an identified autoantigen. For instance, patients with autoimmune demyelinating diseases harboring anti-MOG or anti-AQP4 IgG, could benefit from a rising pool of MOG or AQP4-specific regulatory T cells as in these diseases anti-MOG or anti-AQP4 autoreactive T cells have proven essential in disease pathogenesis (Spadaro *et al*, 2018).

## Materials and Methods

### Animals

A therapeutic protocol of antigen-specific tolerization of 90 days was designed with 6 adults male and female cynomolgus macaques and a preventive protocol of 120 days with 2 adults male cynomolgus macaques, all from the MIRCen colony, imported from a licensed primate breeding center on Mauritius (Cynologics Ltd, Port Louis, Mauritius). Animals were randomly distributed in experimental groups. Following European directive 2010/63/UE and French regulations the project was performed in an agreed user establishment (agreement number 92-032-02), with an institutional permission obtained from the French Ministry of Agriculture after evaluation by an ethical committee (*2015081710528804vl*). All procedures were performed in compliance with CEA’s animal welfare structure. Monkeys remained under veterinary care during the study. Before sample collections, immunization or treatment, animals were sedated with ketamine hydrochloride (Imalgene, 15mg/kg, intramuscular injection) and xylazine (Rompun 2% 0.5mg/kg, intramuscular injection). Anesthesia was maintained during MRI acquisition with propofol (Propovet, 10mg/kg/h, intravenous infusion on the external saphenous vein). Individual animal data are listed in (**supplementary table 1**).

### Study Design and Power Analysis of Test Group Size

In previous experiments we observed that the incidence of EAE in adult cynomolgus macaques of either sex is 95.45% as 21 out of 22 animals developed the disease following immunization with rhMOG/IFA *(unpublished results)*. This indicates that each animal has a theoretical probability of 0.9545 to develop EAE under our protocol. For ethical motives and on behalf of the 3 R principles (Replacement, Reduction and Refinement), we calculated the smallest possible sample of macaques to prove the concept of therapeutic effectiveness of a molecule of which biological significance had been previously measured over 12 macaques (Li *et al*, 2012). We used the Fisher Exact test to calculate the smallest possible sample to have no animal developing the disease. As an acceptable level of confidence of 99% leads to a n = 3, we decided to perform this preclinical study in 2 groups of 3 animals, either treated with anti-DC-ASGPR-MOG or anti-DC-ASGPR-PSA. In a successive experiment to check if intradermal injection of anti-DC-ASGPR-MOG induces anti-MOG IgG, only 2 animals were used. These were also exploited to assess if pretreatment with anti-DC-ASGPR-MOG prevents rhMOG/IFA-induced EAE.

### Immunization and treatment

Animals were immunized every 4 weeks with 300µg of rhMOG (1mg/ml) in IFA (Sigma Aldrich) until disease onset, in the dorsal skin by 6 intradermal injections of 100µl (50µg rhMOG per injection site).

Six animals received subcutaneous injections of 250µg of anti-DC-ASGPR-MOG (treated group) or anti-DC-ASGPR-PSA (control group) every week for 3 weeks starting at the 1st week after initial immunization with rhMOG/IFA and then every first week after each boost with rhMOG/IFA. Each animal received a subcutaneous dose of proteins into the back between the shoulder blades as 5 injections of 100µl of fusion proteins (50µg protein per injection site). In a preventive scheme, 2 macaques received 3 administrations of anti-DC-ASGPR-MOG every week for 3 weeks. They were then immunized with rhMOG/IFA on the fourth week and then again 4 weeks and 8 weeks later. All animals were followed for 90 days after 1^st^immunization with rhMOG.

To assess the phenotype of skin myeloid cells engulfing injected rhMOG, the anti-DC-ASGPR-MOG or a chimeric anti-human CD40-MOG and human IgG4 heavy chain as described above and in (Yin *et al*, 2016), the 3 proteins were conjugated to Alexa Fluorochrome AF488 or AF594 using a microscale labeling kit (Life technology). Ten μg of each protein in 100 μl of PBS was injected intradermally in 2 sites in the back of adult cynomolgus macaques. Skin biopsies (punch 8 mm) were surgically removed at 4 and 24 hrs after injections. Tissues were fixed with 4% PFA in PBS 1x for 6 hrs, dehydrated in 30% sucrose PBS at 4°C, embedded in OCT and frozen at - 42°C. Cryostat sections of 10µm were incubated with further stained with anti-CD163 (GHI-61, 333602 Biolegend), anti-CD68 (KP1, 344716 Dako) or anti-CD1a (5C3, M3571 Dako) overnight at 4°C. A goat anti-mouse IgG1 conjugated to AF594 was used as a secondary antibody. Isotype antibodies were used as negative controls. Tissues were examined using a confocal SP8 microscope (Leica, Germany).

### Clinical observations

Monkeys were observed on daily basis throughout our experiments. Clinical scores were assessed using a semi-quantitative functional scale with severity of disease implying closer endpoints of experiments (Haanstra *et al*, 2013) (**tab. S2)**.

### Fluids collection and immunological analysis

Blood was collected each week, at EAE onset and euthanasia, with a total volume of up to 26 ml per month. Cerebrospinal fluid (CSF) was sampled at EAE onset and euthanasia, for up to 500 µl per puncture. Full hematology with blood cell count (CBC) was performed at each time of bleeding using an HMX A/L (Beckman Coulter). Immunological investigations were performed with fresh whole blood or isolated peripheral blood mononuclear cells (PBMCs).

### T and B cell subsets in blood

Immuno-phenotyping was performed on fresh blood at specified time points by flow cytometry using the following antibodies: anti-CD45 (clone D058-1283, Becton-Dickinson (BD) Le Pont-de-Claix, France); anti-CD3 (SP34-2, BD); anti-CD4 (L200, BD); anti-CD8 (BWl35/80, Miltenyi Biotec, Paris, France); anti-CD95 (DX2, BD); anti-CD28 (clone 28.2, Beckman Coulter); anti-CD69 (FN50, BD); anti-HLA-DR (clone L243, BD); anti-CD20 (2H7, BD); anti-CD27 (M-T271, Miltenyi); anti-IgD (rabbit polyclonal, BioRad, Marnes-la-Coquette, France). Briefly, blood was incubated with antibodies mix for 15 minutes, red blood cells were lysed and cells were washed and fixed. A total event of 10^5^cells was acquired with a BD LSRII analyzer and data analyzed using FlowJo software (Ashland, OR, USA).

### MOG-specific regulatory T cells

Macaque PBMCs (2×10^6^cells/well) were cultured for 44 hr (37°C, 5% CO2) in 500 μl of IMDM (ThermoFisher) supplemented with 10% FCS and 1% penicillin/streptomycin in the presence or absence of 20 μg/ml of rhMOG. Four hours before the end of incubation, Golgi plug (1 µL/mL, BD Biosciences,) was added to media and “Golgi stop” (0.67 µL/mL, BD Biosciences) in each well and cultures were left for another 4 hrs incubation. Cells were washed and stained to detect MOG-specific CD4+ T cell subsets as previously described (Seddiki *et al*, 2014; Brezar *et al*, 2015), using commercial mAbs according to guidelines: anti-CD3-BV768 (SP34-2, BD), anti-CD4-BV605 (L200, BD), anti-CD8-APC Cy7 (SK1, Biolegend), anti-OX40-PE (L106, BD), anti-CD25-BV711 (2A3, BD), anti-CD39-PE-CF594 (TU66, BD) and anti-FOXP3-APC (236A/E7, BD). Intracellular staining for FOXP3 required permeabilization buffer & FOXP3 buffer kit (BD) used as in instructions. Cells were analyzed on a 3-laser LSR II flow cytometer (BD) with at least 10^5^events collected. FlowJo software was used for analysis.

### IgG and IgM antibodies

Plasma anti-rhMOG antibody concentration were assessed by ELISA in 96-well plates. Flat bottom plastic plates (Costar 3595, Corning) were coated with rhMOG (5µg/ml PBS1x) during overnight incubation at 4°C. After washing and blocking with PBS/1% BSA, the wells were incubated in duplicate with 1:200 or 1:2000 diluted plasma samples. Bound cynomolgus monkey antibodies were detected with alkaline phosphate-labeled goat-anti-human IgG (1:1000, 4HI1305, Invitrogen Life Technologies, Bleiswijk, Netherlands) or alkaline phosphate-labeled goat-anti-human IgM (1:2000, A9794, Sigma, St. Quentin Fallavier France). Conjugate binding was quantified with SIGMAFAST p-nitrophenyl phosphate (Sigma, St. Quentin Fallavier France). Measured optical density was converted to arbitrary units (AU) using a concentration curve of the same positive control as reference.

### Cell-based assay (CBA) for titration of antibodies to MOG on living cells

HEK293A cells transfected with the pIRES2-DsRed2-human MOG (HEK293A_MOG) (Horellou *et al*, 2015) were used to detect plasma antibodies binding conformationally intact MOG by flow cytometry. As a control, non-transfected HEK293A cells were used. Briefly, 150,000 cells were incubated with plasma at a 1:50 dilution of plasma for 1 hr at 4°C. Cells were incubated with AF488 goat anti-human IgG1 (1:500, A10631, Invitrogen) for 30 min at 4°C. Cells were fixed in 2% formaldehyde-PBS. A total of 50,000 events per sample were recorded on a FACSCanto II instrument and data analyzed using FlowJo software. Binding was assessed measuring mean fluorescence intensity (MFI). Levels of Arbitrary Units (AU) of anti-MOG antibodies were expressed as *Δ*MFI determined by the subtraction of MFI obtained with control HEK293A from that obtained with HEK293MOG+ cells.

### Cytokine level in plasma

15 cytokines were measured in plasma with multiplex technology, MILLIPLEX MAP human Cytokine Magnetic Bead Panel – Customized Premixed 13 Plex (Bulk) Packaging (IL-1β, GM-CSF, G-CSF, IL-2, IL-4, IL-6, IL-8, IL-10, IL-12p40, IL-17A, TNFα, IFNγ) and MILLIPLEX MAP TGFß Magnetic Bead 3 Plex Kit, (TGFβ1, TGFβ2 and TGFβ3), as 2 separate dosages, following supplier (Merckmillipore, Burlington, MA, USA) guidelines using a Bioplex 200 (BioRad, Hercules, CA, USA). The quantification of all samples was performed all together.

### MRI Acquisitions

Magnetic Resonance Imaging (MRI) on anesthetized animals maintained in a stereotaxic frame were performed on a horizontal 7T Agilent scanner (Palo Alto, CA, USA), using a surface coil for transmission and reception (RAPID Biomedical GmbH, Rimpar, Germany). Lesion segmentation was performed using a high-resolution 2D fast spin-echo sequence. Lesions were manually delineated slice-by-slice by a single operator in order to measure their volume.

### Histology

After euthanasia (Pentobarbital IV, 180 mg/kg) animals were perfused with 2L of ice cold 4% PFA in PBS. Organs including brain, spinal cord, optic nerves, liver, lung, heart, spleen, kidneys, mesenteric and mediastinal ganglia as well as injection and immunization sites were examined and fixed in 4% PFA for 72 hours. All tissues were processed to paraffin blocks, cut and stained with hematoxylin eosin stain (HE). Brain and spinal cord section were stained with luxol-fast-blue (LFB) special stain to assess demyelination. Brain and spinal cord lesions were scored according to their overall severity, number, size, and intralesional myelin loss.

### Immunohistology

Tissues were dewaxed in xylene and rehydrated. Endogenous peroxidase was suppressed by 0.5% H2O2 in methanol. Sections were incubated with rabbit anti-human-IgG (1:100, Sigma, SAB3701291, IgG) and with rabbit anti-human IgM (1:250, Dako). Incubation with primary antibodies was followed by a biotinylated goat-anti-rabbit antibody for 30 min followed by the avidin–biotin–peroxidase complex (Vectastain Elite ABC Kit, Vector Laboratories, PK 6100; Burlingame, CA, USA) for 30 min at room temperature. Positive antigen–antibody reactions were visualized by incubation with 3,3′-diaminobenzidine-tetrahydrochloride (DAB)–H_2_O_2_ in 0.1M imidazole, pH 7.1 for 5 min, followed by slight counterstaining with HE.

### Statistical Analysis

Statistical analyses and graphical representations were done using Prism 5 (GraphPad Software, Inc). Student’s t-test (unpaired, two-sided) was used to compare two groups of values. The two-sided one-way ANOVA test with Tukey’s multiple comparison test was used to compare 3 groups or more values. Heatmaps were generated using R software (R Foundation for Statistical Computing, Vienna, Austria). Hierarchical clustering represented by dendrograms were generated based on the Euclidian distance and using the complete linkage method.

## Acknowledgments

Authors are grateful to Dr Krista G. Haanstra from BPRC in Rijswijk, the Netherlands for her help in setting the EAE model in cynomolgus macaques at MIRCen, CEA, Fontenay-aux-Roses. Authors are grateful to grant sponsors: Infrastructures Nationales en Biologie et Santé (INBS) - 2011 Infectious Disease Models and Innovative Therapies (IDMIT); Grant number: ANR-11-INBS-0008. Baylor Scott and White HealthCare System funding as well as Roche Research Collaborative grants to the Baylor Institute for Immunology Research supported production of the anti-DC-ASGPR-MOG and anti-DC-ASGPR-PSA products. Nicolas Tchitchek was supported by fellowships from the ANRS (France Recherche Nord & Sud Sida-hiv Hépatites).

## Author contributions

AH, CC, CL, CMF, CS, JF, JM, LS, NS, NT, PhH, ST, VC: contributed to the acquisition and analysis of data. CMF, CS, JF, JM, VC: Animals follow-up, samples collection and MRI. CMF, CS, JM, VC: laboratory dosages. AH, CL, NS, PhH: cytometry. CC, LS, ST: tissues treatments and histology. CMF, CS, LS, NT, VC: graphs and statistical analysis. B’tH, CMF, CS, GZ, KD, LS, NS, NT, PhH, PK, RLG, SO, SZ: manuscript drafting for main intellectual content. All authors gave final approval to the article.

## Data and materials availability

All data associated with this study are available in the main text or the supplementary materials.

## The paper explained

### Problem

Brain autoimmune diseases are a major concern of public health and the first cause of demyelinating processes in children and adults. These conditions are currently treated with immunosuppression while awaiting cures able to restore normal immune response and health. Antigen-presenting cells (APC) are specialized immune cells sited in all tissues with remarkable capacity to direct adaptive lymphocyte response toward immunity or tolerance. Thus, manipulation of APC offers a prospect to boost regulatory lymphocytes to inhibit damaging immune response, especially in autoimmune processes with an identified autoantigen as in the case of relapsing encephalitis coursing with antibodies to the Myelin Oligodendrocyte Protein (MOG).

## Results

We investigated the effect of a vaccination procedure to raise immune tolerance to the antigen MOG in a macaque model of experimental autoimmune encephalitis (EAE) in which an autoimmune response mediated by MOG-specific autoreactive lymphocytes and anti-MOG antibodies, elicits brain inflammation and demyelination. To target MOG to tolerogenic APC, we used a recombinant antibody binding to the Dendritic Cells (DC)-Asialoglycoprotein receptor (DC-ASGPR). The skin injection of an anti-DC-ASGPR-MOG fusion protein, but not a control anti-DC-ASGPR-PSA (prostate specific antigen) protein, remarkably prevented the burst of the autoimmune encephalitis in monkeys. The treatment with the anti-DC-ASGPR-MOG was associated to a rise of MOG-specific regulatory lymphocytes, which prevented lymphocyte activation and the production of pro-inflammatory cytokines seen in control animals developing an autoimmune encephalitis. Moreover, a significant upsurge of the anti-inflammatory cytokine TGF*β* was observed after MOG re-immunization in animals receiving the effective treatment, specifying the protective response elicited by the induced MOG-specific regulatory cells.

## Impact

Our experiments indicate that a vaccination scheme for immune tolerization towards a self-antigen is highly efficient in preventing autoimmune aggression in primates. This is notorious as contrary to mouse strains, macaques are genetically diverse, indicating that the vaccine efficiency is not restricted to a specific genotype and is the result of a general mechanism of immune regulation. The results of this preclinical protocol raise great hopes for the treatment of autoimmune conditions in humans through targeting of self-antigens to skin APC expressing DC-ASGPR. Such treatment promises to be of easy administration, safe and highly efficient.

