## Supplementary data for "Targeting MOG to skin macrophages prevents EAE in macaques through TGFβ-induced peripheral tolerance"

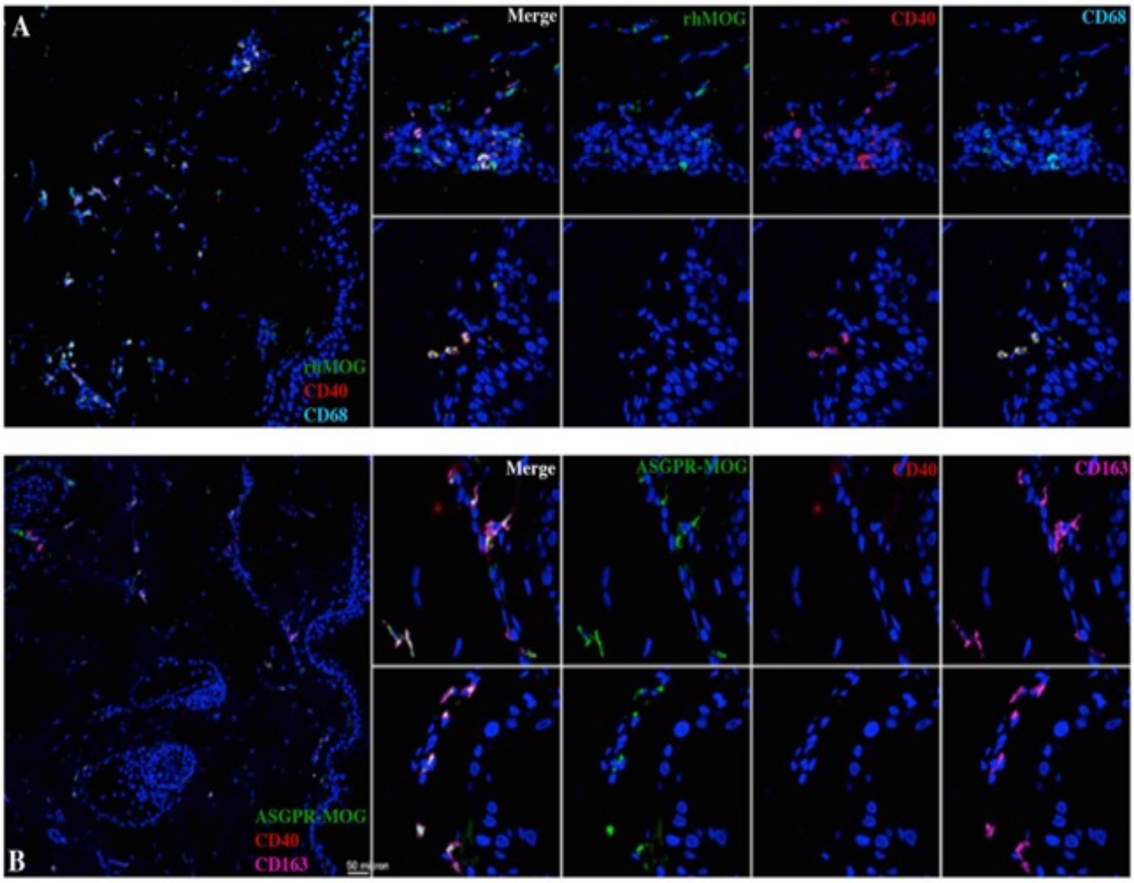


**Figure S1. Recombinant MOG or anti-DC-ASGPR-MOG recombinant proteins entry in dermal myeloid cells.** Through IHC of NHP skin we observed, **(A)** that injected rhMOG-AF488 protein in green and the anti-CD40-MOG-AF594 antibody (red) colocalize in DC or macrophages expressing CD68 (cyan) as well as in Langerhans cells close to the epidermis expressing CD1a (yellow). **(B)** The anti-DC-ASGPR-MOG-AF488 (green) was detected in dermal macrophages expressing CD163 (magenta) but did not colocalize with the anti-CD40-MOG (red). Cells nuclei were stained with DAPI (blue).

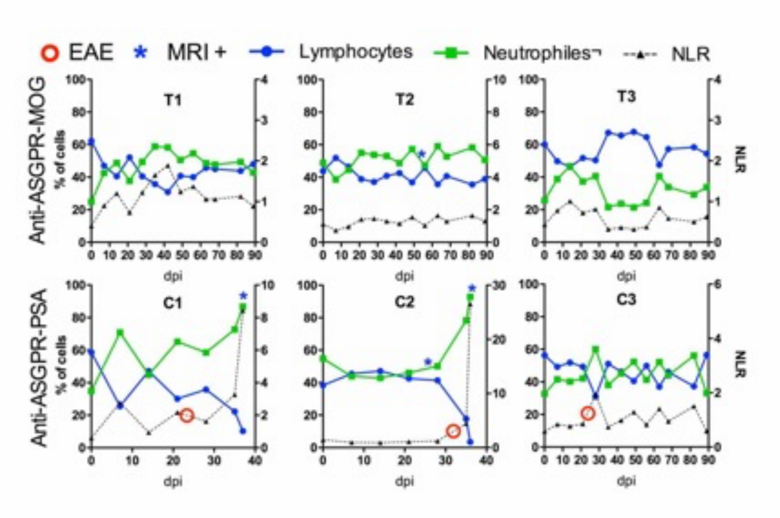


**Figure S2. Onset of EAE and neutrophils/lymphocyte ratio**. Levels of circulating lymphocytes in blue circles and neutrophils in green squares during the experiment in animals immunized with rhMOG/IFA and then treated with anti-DC-ASGPR-MOG (T1, T2, T3) or with anti-DC-ASGPR-PSA (C1, C2, C3). The doted lines represent les neutrophils/lymphocyte ratio (NLR), which increases few days before EAE onset. Red circle: EAE onset; blue star: positive MRI displaying an hyperintense image; dpi: days post immunization.

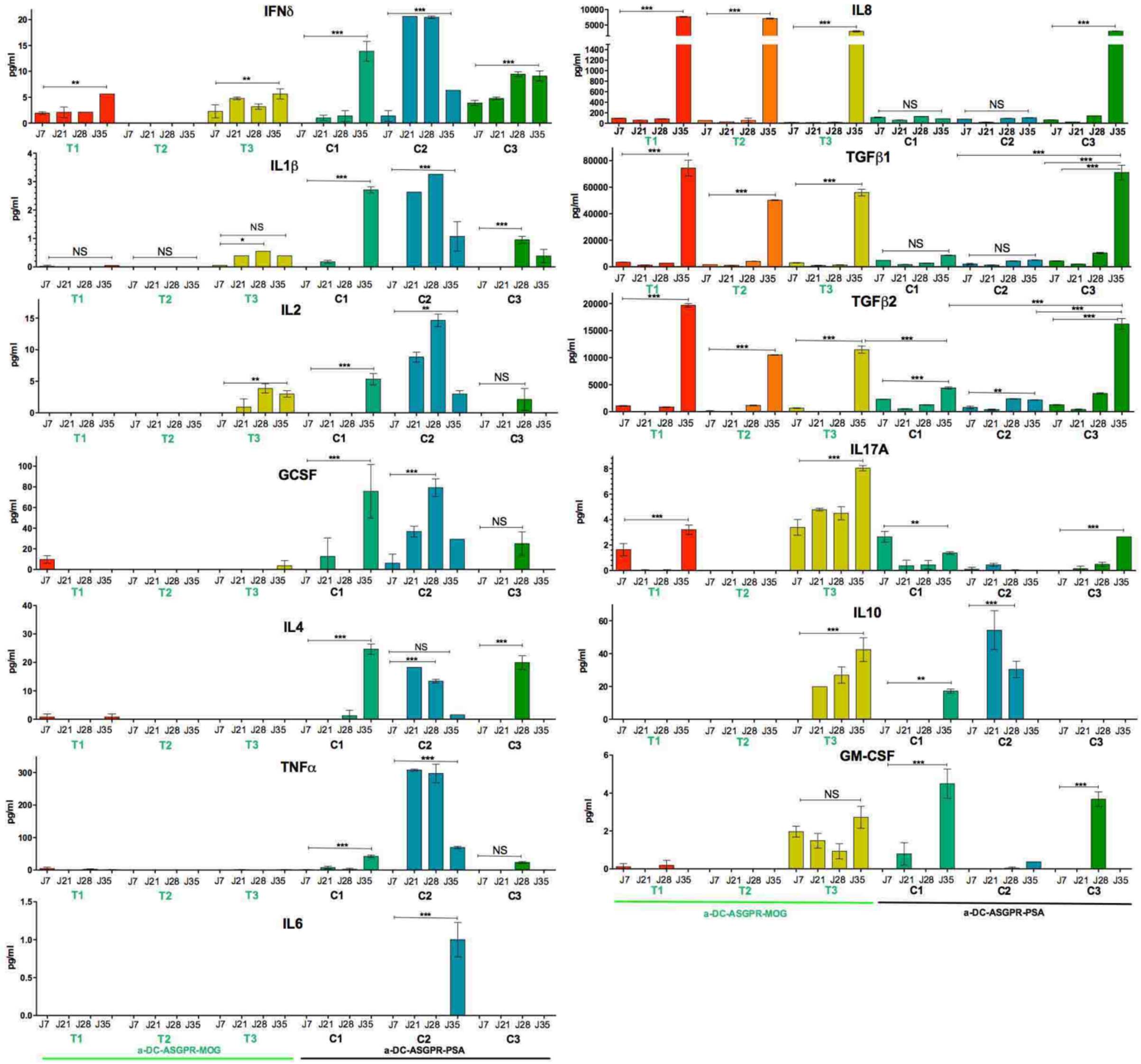


**Figure S3. Levels of cytokines per animal at different time points.** Each bar rates the concentration (pg/ml) of a particular cytokine in plasma of animals treated with anti-DC-ASGPR-MOG (T1, T2, T3), underlined in green and control animals (C1, C2, C3) underlined in black. Graphs stacked on the left side represent cytokines mostly expressed in control animals. Graphs on the right display cytokines mainly expressed in treated animals. Bars represent mean values of 2 dosages ± SEM. Statistics, one-way ANOVA and Tukey posttest; $p > 0.05$; (*) $p < 0.05$; (**) $p < 0.01$; (***) $p < 0.001$.

| Animals | Age (y) | Weight (kg) | Sex | Immunization | Treatment |
|---|---|---|---|---|---|
| T1 | 9 | 8,3 | M | rhMOG/IFA 300µg | Anti-DC-ASGPR-MOG |
| T2 | 6 | 4,3 | F | | |
| T3 | 7 | 3,8 | F | | |
| C1 | 10 | 6,2 | M | | Anti-DC-ASGPR-PSA |
| C2 | 6 | 3,8 | F | | |
| C3 | 10 | 4,6 | F | | |
| P1 | 8 | 9,58 | M | Anti-DC-ASGPR-MOG | rhMOG/IFA 300µg |
| P2 | 8 | 8,75 | M | | |

**Table S1.** Characteristics of animals involved in this protocol and their respective conditioning and treatment. Animals immunized with rhMOG and then treated with anti-DC-ASGPR-MOG (T); Control animals immunized with rhMOG and then treated with anti-DC-ASGPR-PSA (C); Animals preventively vaccinated with DC-ASGPR-MOG and then immunized with (P).

| EAE score | Clinical signs | Maximum duration |
|---|---|---|
| 0 | No clinical signs. | End of experiment |
| 1 | Behavioral discomforts without effect on social behavior and feeding (tremors, scratching, stereotypes, nystagmus, eyelid ptosis, paresis with persistent limb using). | 20 weeks |
| 2 | Behavioral discomfort with an impact on feeding or social behavior (tremors with dysmetria, oculomotricity paralysis, paresis with underusing of one limb and counterbalanced by other limbs). | 4 weeks |
| 3 | Motor impairment with an impact on the function but not on feeding or social behavior (lameness, ataxia, imbalance). | 2 weeks |
| 4 | Behavioral and self-feeding impairments (paralysis, paresis). | < 18h |
| 5 | Coma (breathing normally and responding to external stimuli). | < 6h |
| 6 | Moribund, no response. | < 1h |

**Table S2.** Clinical score attribution to behavioral and neurological deficit in the course of EAE. The higher the score the more sever is EAE and the shorter is the accepted time of survival.

| Histopathological characteristic of the lesions | Controls | Treated |
|---|---|---|
| Overall severity | Severe | Mild |
| Number and size | 2-3 large lesions (> 1cm in diameter) | 1 small lesions (< 0.5 cm in diameter) |
| Principal characteristic | Necrosis and hemorrhage of the white matter | Demyelination without necrosis |
| **Principal infiltrating cell type** | | |
| • Neuropil (WM) | Neutrophils, Macrophages | Macrophages |
| • Perivascular space | Mixed (neutrophils, macrophages, +/- lymphocytes) | No infiltration or mononuclear (macrophages, lymphocytes) |
| Myelin phagocytosis | Present | Low or absent |

**Table S3.** Histological characteristics observed in lesions of an animals treated with anti-DC-ASGPR-MOG (Treated) as compared to controls treated with the anti-DC-ASGPR-PSA antibody (Controls).

| | Day 0 | Day 7 | | Day 21 | | Day 28 | | Day 35 | |
|---|---|---|---|---|---|---|---|---|---|
| | NAIVES | ASGPR MOG | ASGPR PSA | ASGPR MOG | ASGPR PSA | ASGPR MOG | ASGPR PSA | ASGPR MOG | ASGPR PSA |
| **Group 2 (EAE incubation)** | | **T1, T2, T3** | **C1, C2, C3** | | **C1, C3** | **T1, T2** | **C1** | | |
| **Group 4 (EAE onset & progression)** | | | | | **C2** | | **C2, C3** | | **C1, C2** |
| **Group 1 (EAE solving)** | | | | **T1, T2, T3** | | **T3** | | | |
| **Group 3 (EAE resolution)** | **N1, N2, N3, N4** | | | | | | | **T1, T2, T3** | **C3** |

**Table S4.** Tabular representation of the heatmap presented in **(figure 5)** ranking animals by cytokine levels at different time points (days post-immunization with rhMOG/IFA). From right to left the heat map segregates 4 groups, from low to high amounts of cytokines and numbered from 1 to 4. In this table we interpret each group as representing a particular step of EAE from incubation (upper row) to onset and progression (second row), to EAE solving (3$^{rd}$ row) and complete resolution (bottom row). In the upper row, group 2 associates treated and control animals at several time points including 7, 21 and 28 dpi. As it excludes naïve animals, it likely represents a cytokine picture corresponding to disease incubation. In the second row, group 4 includes all highest amounts of pro-inflammatory cytokines in control animals treated with anti-DC-ASGPR-PSA at EAE onset and at peak of disease. In the 3$^{rd}$ row, as group 1 associates only animals treated with anti-DC-ASGPR-MOG at 21 or 28 dpi with a particular pattern of low levels of TGFβ and high levels of IL-10, it seems to correspond to a phase of immunomodulation for disease solving induced by the treatment with anti-DC-ASGPR-MOG. In the bottom row, group 3 associates only animals not developing EAE at 35 dpi and naive animals; it thus corresponds to a cytokine picture of disease resolution and immune homeostasis with the highest levels of TGFβ.
